# A low-cost and easy-to-use cell preservation reagent for 4°C or room temperature sample storage

**DOI:** 10.1101/745232

**Authors:** Sidharth Shankar, Shreya Roy, Ari Geary-Teeter, George M. Martin, Paula D. Ladd

## Abstract

Current biospecimen storage and preservation methods no longer meet the demands of basic research and clinical diagnostics. Biospecimen preservation methodology has not advanced to accommodate cutting edge molecular analysis technologies that target single cells and full-length transcripts. Traditional methods, such as flash freezing and formalin-fixed paraffin-embedding (FFPE), were designed to provide information on cell structure and spatial relationships in whole tissues. These methods, however, do not maintain the integrity of proteins and nucleic acids. In this proof-of-concept study we examine preservation mechanisms utilized in nature for survival during cold seasons or periods of drought. Plants, brine shrimp, and tardigrades rely on components such as disaccharides or intrinsically disordered proteins to maintain cellular structure and biological activity in the settings of these environmental stresses. Our study demonstrates that these reagents aid in mammalian cell preservation for at least one week when a sample is stored in solution at 4°C, or as a dried sample in a low humidity desiccator.

## Introduction

Quality and reliable tissue sample preservation is of paramount importance in basic biological and clinical research, as well as for diagnostic and core life science service laboratories. Variations in preservation methodology impact downstream assays, which is of particularly concern in clinical pathology and diagnosis (1–3). Major considerations include delays in preservation, length of time the sample exists in preservation reagents, and improper storage conditions (4). The most common preservation methods, FFPE and flash freezing samples, while in wide use, have significant limitations (5).

FFPE is a widely adopted preservation method to preserve proteins and tissue structure and is nearly ubiquitous in anatomic pathology laboratories. The basic protocol involves fixing a tissue sample in formalin and then embedding the sample in paraffin wax to facilitate dissection into thin tissue sections for placement on microscope slides (6). At this point the section can be stained with reagents to permit highly reliable diagnoses of a wide range of pathologies. An advantage of FFPE is that tissue structure is well preserved, and samples can be stored at room temperature for many years. Unfortunately, this method requires toxic chemicals, specialized training, and is sensitive to individual processing differences. Moreover, a disadvantage of this method is that it results in denatured, non-biologically active proteins and degraded genomic DNA and RNA and thus does not address the growing molecular biological advances for more precise clinical and research characterizations of pathologic processes. Formalin-free methodologies, such as the PAXgene Tissue system (PreAnalytiX, Switzerland) exhibit improved nucleic acid quality, but only when samples are stored at −20°C (7). Storage of PAXgene preserved samples at room temperature is associated with nucleic acid degradation, much like with FFPE.

Cryopreservation is another widely used preservation method; samples are stored in −80°C freezers or liquid nitrogen (−196°C). At these temperatures, metabolic activity is virtually halted, and cells can remain stable for decades as long as low temperatures are maintained. An advantage of freezing a sample is that proteins can be biologically active when thawed. Moreover, cryopreservation requires little training. As for the case of FFPE, however, delays in freezing can induce considerable variability in preservation among samples (8). A major disadvantage of this method is a potentially crippling reliance on electricity or liquid nitrogen, placing samples at high risk during periods of natural disaster (9). Furthermore, cell membranes are typically damaged in the freeze thaw process, hindering single cell isolation.

Cryoprotectants are often used to ameliorate cellular damage during the freezing and thawing processes (10). DMSO and glycerol are typical cryoprotectants when cells will be returned to a state of growth. A reagent designed to enhance RNA stability is RNAlater, which includes a solution of saturated ammonium sulfate, EDTA, and a sodium citrate buffer at a pH of 5.2. The ammonium sulfate denatures the proteins in the sample, including nucleases and RNA binding proteins, which then can remain wrapped around RNA to protect it from exogenous RNases. An advantage here is that samples can be stored in this solution for short periods of time (one week) at room temperature, up to one month at 4°C, and indefinitely at −20°C. The disadvantage of this method is that, similar to that of FFPE and freezing, cell membranes and global proteins are denatured, hindering single cell isolation. Moreover, numerous users have reported reduced RNA yields and global transcript biases (11–13).

Given the disadvantages of FFPE, freezing, and RNAlater, it is appropriate to consider alternative methodologies that are not only compatible with molecular analysis, but also are easy to use and relatively non-toxic. In search of innovative approaches to the development of such improved methodologies, our group reviewed the literature on Nature’s examples of cryptobiosis, a reversible pausing of the physiological state used by a wide range of organisms, including microbes, plants and animals (14). More specifically we were interested in a cryptobiosis subtype, anhydrobiosis (extreme drying). During anhydrobiosis a protective “glass matrix” is formed to protect the organism.

One common adaptation essential to some desiccation tolerant organisms is the use of sugars, specifically disaccharide sugars, sucrose and trehalose. These sugars prevent protein denaturation and protect against membrane degradation during a dehydrated state, possibly by water exclusion or water replacement (15). However, not all desiccation tolerant organisms express trehalose. Instead, some tardigrade species utilize intrinsically disordered proteins (IDP) to protect the organism during extremely dry conditions (16). Tardigrade specific IDPs form cytoprotective crystalline amorphous solids during desiccation and abiotic stress tolerance (17).

To test our hypothesis that mammalian cells could be preserved in a dry environment, we selected trehalose and bovine serum albumin (BSA), which contains several regions with flexible disordered domains (18). Here, we demonstrate that we can isolate high quality DNA and RNA from human diploid fibroblast cells preserved in PBS containing either trehalose or BSA and supplemented with a chelating agent, EDTA. We compared cells preserved for one week in solution at 4C or desiccated at room temperature to cells that had been frozen for the same time periods, as well as freshly harvested samples. Under those conditions, our reagent is capable of preserving a sample without significant macromolecular loss and is very easy to use. We posit that this reagent could serve as a novel, relatively nontoxic and widely accessible means of protecting cellular components that are likely to become of crucial importance as pathologists seek to apply newly available tools resulting from advances in molecular biological research.

When growing tissue culture cells, it is common to store cell pellets in a −80°C freezer. These pellets are often used to represent the untreated population or time zero control samples. Typically, adherent cells are enzymatically released from the culture dish, centrifuged to form a cell pellet, and stored in the freezer. The potential for variation is high in this multistep process; prolonged enzymatic treatment alters gene expression and high centrifugal force damages cell membranes. Additionally, freezing induces ice crystals which break cell walls and long-term storage at −80°C is known to fragment both DNA and RNA (19).

### Preservation of Nucleic Acids in Solution

To determine whether high quality nucleic acids could be isolated from tissue culture cells stored at 4°C instead of −80°C, we established a simple protocol to preserve tissue culture cells. First, the growth media is removed from dishes of human diploid fibroblast cells. The cells are rinsed two times with PBS, and the final wash buffer is completely aspirated (Figure 1A left most panels). One of the four preservation reagents, or RNAlater, is added to each dish and then the dish is stored in a refrigerator at 4°C (Diagram 1A middle panels). A comparably plated dish is lysed to represent the RNA present at the time of preservation. After one week, all dishes stored at 4°C are lysed, and RNA is extracted. This protocol relies on common tissue culture techniques, namely pipetting, vacuum removal of media and buffers, and thus can be easily performed by all laboratory personnel. Furthermore, this protocol does not require enzymatic treatment and the original growth dish can be used, reducing labelling errors.

**Figure 1.**
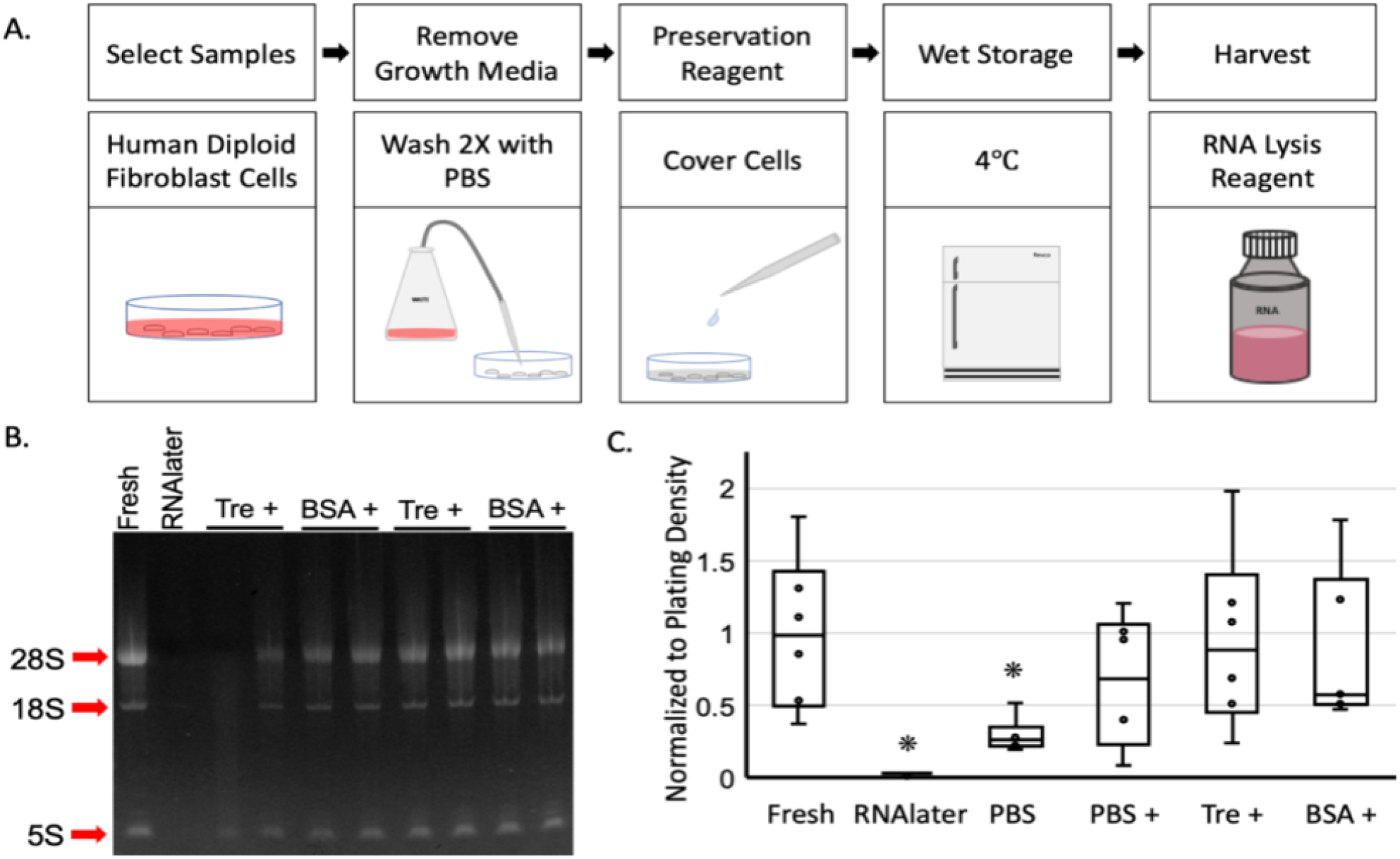
High quality RNA can be extracted from cells stored in a solution containing EDTA at 4°C for one week. A) Cartoon of protocol followed to preserved cells in solution. B) Representative 1% agarose gel with RNA from freshly harvested cells and compared with RNA from cells stored in RNAlater, PBS, PBS +, Tre +, and BSA +. Red arrows denote 28S, 18S, and 5S. C) Box plot depicting range of RNA concentrations, center bar indicates median. * denotes p < 0.02 N = 6

Four preservation buffers were selected based on our crowdfunded project titled: *Developing a new tissue preservation method inspired by Tardigrades* (20). All four preservation reagents contain phosphate buffered saline (PBS). Three of the reagents are PBS supplemented with EDTA, a chelating agent that sequesters metal ions and reduces enzymatic activity of metal ion-dependent enzymes (21). One of the PBS plus EDTA reagents also contains Trehalose (Tre), a disaccharide found in many desiccation tolerant organisms, while another contains molecular-grade bovine serum albumin (BSA), a protein with multiple intrinsically disordered domains (22, 23). The last reagent is PBS solely supplemented with EDTA. For the remainder of this paper, the four reagents described above are indicated as PBS, PBS+, Tre+, and BSA+, the plus sign indicating the presence of EDTA.

## Results

The preservation capability of each of the four reagents, PBS, PBS+, Tre+, BSA+, as well as a commercial reagent, RNAlater (Ambion Thermo Fisher, Waltham, MA), were compared to RNA isolated from freshly harvested cells to determine how well RNA is preserved for one week at 4°C. Global RNA integrity is similar for the freshly isolated RNA sample and RNA that was preserved for one week at 4°C with each of the reagents, PBS+, Tre+, and BSA+ (Figure 1B). Visually, we can see 2-3 times more of the 28S rRNA relative to 18S in the fresh sample as well in the samples supplemented with EDTA. A high 28S/18S ratio indicates high quality RNA that has not degraded.

The samples preserved with PBS alone were either degraded or faintly present (Figure 1B, PBS lanes and supplemental Figure 1A and C). All of the samples preserved with RNAlater contained only trace levels of RNA (Figure 1B, RNAlater lanes and supplemental Figure 1A and C). Initially, this was a surprising result, as the RNAlater reagent had been used by our team multiple times several months prior to these studies with far better output. This suggests that the RNAlater reagent has a shorter shelf life than the PBS-based reagents.

The RNA concentration was similar for all of the samples preserved for one week at 4°C in PBS+, Tre+, and BSA+ (Figure 1C). No significant difference was apparent relative to freshly isolated RNA from equally plated dishes. In this data set, the average RNA concentration from the Tre+ and BSA+ preserved cells were 94% and 84% of freshly isolated RNA, respectively, while PBS+ samples were 66%. These results are in sharp contrast to the PBS samples, which had a concentration that was less than 33% of the fresh samples. This suggests that EDTA is a key component in the preservation of RNA integrity and concentration. Adding trehalose or BSA appears to have some impact on total RNA concentration; however, there is no significant difference between the three preservation reagents. We can conclude that adding trehalose or BSA does not have a negative effect on RNA content.

We also examined whether genomic DNA (gDNA) could also be isolated from the samples stored wet at 4°C. Here we demonstrate that the DNA integrity is similar for freshly isolated samples and samples stored frozen or wet at 4°C for one week (Supplemental Figure 2A). Unfortunately, DNA concentration was reduced in all buffer conditions (Supplemental Figure 2B). This reduction was seen in most of the frozen samples, suggesting that DNases are active during the freeze/thaw process.

Overall, this proof-of-concept study demonstrates that cells stored in solution at 4°C for one week in PBS with EDTA and Trehalose is correlated with RNA concentration and RNA integrity similar to RNA freshly isolated from equally plated cell cultures. This has substantial implication for researchers who want to transport cellular material to collaborators or core facilities, as shipping costs are reduced when dry and wet ice is not needed.

### Nucleic Acid Stability after Dry Preservation

To determine whether the same reagents could be used to preserve diploid fibroblast cells at room temperature in a dry format, we modified the wet preservation protocol to include a drying step. After the preservation reagent is added to a dish of cells, the dish is placed under a table fan until dry. A 60 mm dish of cells typically dried in 5 minutes or less, thus requiring little extra effort to preserve a sample (Figure 2A). We used a 9-inch diameter fan, so that all of the dishes could be dried at one time, eliminating drying variation. Once the dishes were dried, the samples were placed into a desiccator with fresh drierite and 30 gram food-grade silica pouches. A humidity indicator card did not register any humidity in the desiccator during storage. As a point of reference, samples with equal cell density were washed twice with PBS and frozen at −80°C for one week. RNA isolated from the dry preserved and frozen samples were compared to RNA isolated from non-preserved cells (Figure 2B and C).

**Figure 2.**
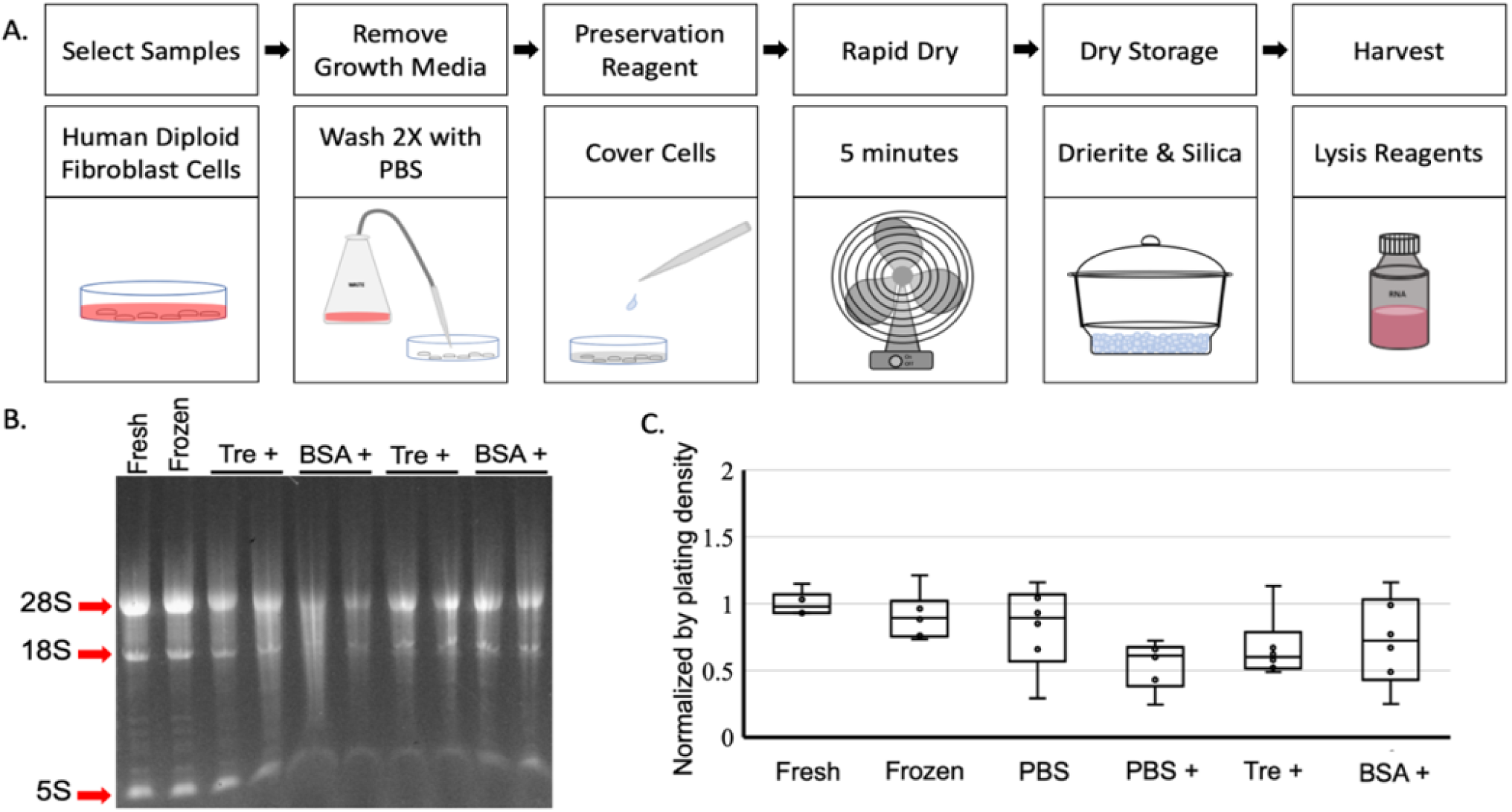
High quality RNA can be extracted from cells stored in a desiccator for one week. A) Cartoon of protocol followed to preserve cells in solution. B) Representative 1% agarose gel with RNA from freshly harvested cells and compared with RNA from cells stored frozen, in PBS, PBS +, Tre +, and BSA +. Red arrows denote 28S, 18S, and 5S. B) Box plot depicting range of RNA concentrations, center bar indicates median. N = 6 * denotes p < 0.015

The RNA isolated from dry preserved samples were similar to RNA from fresh and frozen samples, but for the most part had an additional smear within the lane (Figure 2B and Supplemental Figure 2A and C). In the dry preserved samples, 28S and 18S are visible, but do not resolve appropriately in the agarose gel, as if there is something present in the sample to prevent resolution of 5S RNA (Figure 2B, Supplemental Figure 2A and C). Samples preserved in PBS or PBS+ were the most impacted, though streaking is apparent in some of the Tre+ and BSA+ samples. This streaking is not seen in the RNA samples from fresh or frozen cells, suggesting this is not an artefact of our RNA isolation and rehydration reagents. It is important to highlight that the primary difference in the cell populations from fresh, frozen, or dried preserved samples was the presence of liquid in the cytoplasm. In our studies, we did not hydrate the dry preserved cells prior to lysis as we wanted to first determine if RNA was intact after dry storage. Perhaps this streaking artefact is due to salt interaction with the RNA backbone during preservation in the presence of phosphate buffered saline. RNA sample dilution improves gel resolution, suggesting the artefact is soluble.

In the dry condition, the average RNA concentration from cells preserved in PBS without EDTA is nearly as high as RNA from fresh and frozen samples. This suggests that removing water from a sample is key to preservation and is consistent with results that employ lyophilization of samples. Our protocol requires less training and equipment than lyophilization, a method of freeze-drying for storage at room temperature, thus benefits laboratories with fewer resources (24, 25).

The RNA concentrations for samples preserved in PBS+ or in Tre+ are significantly lower. (Figure 2B, p <= 0.015). This result is consistent for PBS+ preserved samples, but not for the Tre+ sample. This difference in reagent performance in wet and dry preservation conditions could be due to the sugar coating that formed after drying. Since we did not solubilize the sample with water before RNA isolation, the sugar-coating hindered lysis. We noticed that the Tre+ samples were particularly difficult to transfer after adding lysis buffer. We also noticed that the Tre+ samples were particularly difficult to transfer after adding lysis buffer, suggesting that sample rehydration should be examined before concluding that EDTA and or trehalose are not beneficial for dry sample preservation in a desiccator.

Genomic DNA can be isolated from dry preserved samples. In contrast to the RNA samples, there was a loss of gDNA integrity in samples preserved with PBS or BSA+ (Supplemental Figure 3A). However, gDNA concentrations were similar for samples stored dry or in a freezer (Supplemental Figure 3B). While harvesting DNA from fresh samples is the most ideal approach, it is not always an option, such as when collecting field samples. Furthermore, carrying ice and storage coolers into the field can be a challenge. Dry storage is an ideal solution for field scientists, as well as for shipping samples without needing ice.

## Discussion

In this study we have examined sample preservations in two different contexts. In the first condition, the sample is stored in solution at 4°C, while in the second condition, the sample is stored in dry preservation at room temperature and very low humidity.

We have demonstrated, in the first experimental condition, that high quality total RNA can be isolated from human diploid fibroblast cells stored at 4°C in a solution of PBS supplemented with EDTA. Adding trehalose, a disaccharide upregulated in multiple cryptobiosis-tolerant organisms, improves total RNA concentration, though it is not necessary. Furthermore, BSA does not provide any improvement relative to PBS with EDTA. Thus, when samples are held at 4°C, the key component for nucleic acid preservation is EDTA, a metal ion chelator. The most plausible explanation is that EDTA reduces nuclease activity by chelating cofactors, such as Mg^2+^, Cu^+^ and Mn^2+^.

In the second experimental condition, we were able to isolate high quality total RNA from human diploid fibroblast cells that were dried in the presence of the same preservation reagents, and then stored at very low humidity in a desiccator at room temperature. In contrast with the samples stored in solution at 4°C, samples stored in PBS without EDTA had nearly the same concentration of total RNA as fresh and frozen samples. However, the overall RNA quality is impaired in this condition, with apparent nicking of the RNA strands and potentially protein aggregation. Adding trehalose or BSA may mitigate this effect somewhat, but further investigations of drying speed and rehydration conditions are needed to fully understand what will be required for an optimal technique. Nonetheless, it is provocative that a cell sample can be air dried at room temperature and stored in a desiccator for a week. Lyophilisation typically requires freezing the sample to a very low temperature and then drying the sample under vacuum. We have demonstrated that laboratories with few resources could preserve samples in dry form, for at least the short term.

In contrast to total RNA, gDNA protection was not enhanced by the presence of EDTA when the samples are stored in solution at 4°C. This was an unexpected finding, as DNA is generally more stable than RNA. A possible explanation is the fact that not all DNases require Mg^2+^ ions, thus avoiding the inhibitory effects of EDTA. As an example, DNase IIα, a lysosomal enzyme essential for DNA waste removal, does not use a metal ion as a cofactor and remains unaffected by EDTA (26, 27). This enzyme and perhaps others were able to maintain enough activity during the wet preservation process to degrade the gDNA within each sample. It is unclear if this could also explain the reduced gDNA content in several of the frozen samples; perhaps there is more to learn about enzymatic activity during the freeze/thaw process.

When stored dry, gDNA integrity is compromised for samples dried in the presence of PBS only or when BSA is present. Here, we can only speculate whether EDTA is capable of providing greater protection, as demonstrated by an increase in gDNA concentration for PBS+ and Tre+ samples. However, this protection did not extend to the BSA+ samples. Note that in dry conditions, BSA increases the total protein content of the sample. Thus, when lysis reagent is added to the sample, it is plausible that the higher protein content sequesters gDNA, which is removed with the precipitated proteins. This is in contrast to the samples stored in solution at 4°C, as the preservation reagent is removed prior to lysis.

Our dry preservation method does not mimic preservation found in nature. During the onset of cryptobiosis, organisms are sensing and responding to changes in the environment by altering expression of genes and proteins to induce endogenous preservation pathways and additionally degrading transcripts and proteins that are not beneficial. It is equally crucial that organisms have the ability to revert these processes to return to their normal state post-desiccation. Of course, a preservation technique capable of preserving a sample in a suspended state and returning it to normal homeostasis would be a breakthrough in clinical and pathology laboratories, as well for transplantation laboratories. Rather than a permanent halting, a paused state with the potential for future viability could lead to more detailed secondary analysis on patient samples, particularly biopsies, and likely more accurate diagnoses. A first step in this direction is the maintenance of physiological pH in our reagent. Reagents such as RNAlater and DNA/RNA Shield lower the pH, thus denaturing the proteins, and ultimately preserving cellular materials in non-physiological pH. Further, there exist numerous enzymes that are active in acidic environments perpetuated by these reagents. By creating a reagent at a physiological pH, we reduce the activity of these enzymes, and further our goal of halting all enzymatic and metabolic activity, without considerable damage to cellular constituents.

Our study underscores the challenges researchers face when choosing a preservation method during the course of a study, as well as in biospecimen preservation for clinical analysis. In research studies, consistency is key. When all samples are preserved in the same manner, differences between each sample are more convincing. But in certain situations, such as with clinical samples where variation is harder to manage, having a preservation reagent that is easy-to-use and that immediately arrests biological activity is key to specificity and accuracy of a test. Our preservation reagents have the ability to serve dual purposes; they may serve as a stand-alone means of cell preservation or potentially as an enhancing agent in the pre-preservation of clinical samples.

Future studies should move from cultured cells to solid tissues from experimental mammalian materials, ideally using laboratory strains of mice, as this would provide the potential to optimize macromolecular integrity as functions not only of a wide range of tissues but also from animals of different ages and with a variety of pathologies.

## Conclusion

In this proof-of-concept study, we demonstrate that a monolayer of fibroblast cells can be stored in solution for one week at 4°C or dried in the presence of these solutions and stored at room-temperature, as long as the samples are shielded from humidity. Full length DNA and RNA can be extracted from these samples. Humidity is key, as water is a key component of life. Without humidity/water, proteins have no (or minimal) activity. There is no degradation because there is no activity. Our results indicate there could still be protein aggregation, demonstrating that adding trehalose and/or BSA helps to reduce aggregation.

At a time when low input analysis is needed for clinical diagnostics and monitoring, improved biospecimen preservation will enhance the contemporary molecular methodologies that are being used to understand genomics, transcriptomics, epigenomics, proteomics, and metabolomics.

## Materials and Methods

### Cell Culture

Low passage primary human diploid foreskin fibroblast cells were used for preservation experiments. The cells were cultured in DMEM (Gibco, ThermoFisher Scientific) supplemented with 10% fetal bovine serum and 1% PenStrep (100 U/mL Penicillin /10 μg/mL Streptomycin). The cells were maintained under standard cultural conditions at 37°C in an atmosphere of 5% CO_2_ and 5% O_2_ in a Heracell CO2 incubator (ThermoFisher Scientific, Waltham, MA) (28). Gas phase O2 tensions were controlled by continuous injection of appropriate amount of medical grade N2 to reach the target levels of oxygen. Cells were seeded at equal cell densities in 12-well, 6-well, or 60 mm dishes for the isolations of protein, DNA, or RNA.

### Preservation

Reagent #1 is PBS, pH 7.4, the base of all conditions. Reagent #2 is PBS plus 10 mM EDTA. Reagent #3 contains PBS, plus 10 mM EDTA and 50 mM Trehalose (ThermoFisher Scientific, Waltham, MA). Reagent #4A is PBS, plus 10 mM EDTA and 5% fetal bovine serum. Reagent #4B contains PBS, pH 7.4 plus 10 mM EDTA and 5% molecular grade BSA (ThermoFisher Scientific, Waltham, MA).

The volume of preservation reagent was different for each vessel size. 75 μl, 100 μl, and 150 μl of reagents 1-4, were used for 12-well, 6-well, and 60 mm dishes respectively. A 9-inch oscillating fan (SMC, China) was used to rapidly dry each dish of reagent and cells. The dry dishes of cells were immediately placed in a glass desiccator in a dark cabinet. Less than 10% humidity was achieved using drierite anhydrous calcium sulfate (W.A. Hammond Drierite Co Ltd, Xenia, OH) and food grade silica packs (Dry & Dry, Silicagel Factory, Brea, CA).

For each preservation trial, untreated samples of equal cell densities were frozen at −80C for the same duration as the dry preservation samples. Samples for all preservation reagent conditions were harvested at 1 week, 1, 2, and 3 months. Fresh samples of equal cell densities were harvested on day zero to represent the starting levels of protein, DNA, and RNA.

### Nucleic Acids

DNA was extracted using in-house reagents based on the Gentra-Puregene methods. RNA was extracted using Trizol, as per the manufacture’s protocol (Qiagen, Germantown, MD). RNA and DNA concentrations were determined using a Nanodrop Lite Nano Spectrophotometer (ThermoFisher Scientific, Waltham, MA) and visualized on 1% agarose e-gel Power Snap (ThermoFisher Scientific, Waltham, MA).

### Statistical analysis

Statistical significance was determined by the Student t-test.

## Acknowledgements

We wish to acknowledge Dr. Jenny Tenlen, PhD, Biology Department, Seattle Pacific University and Dr. Silvan Urfer, DMV, Department of Pathology, University of Washington for their advice and support of this project. We also want to acknowledge all project backers on Experiment.com/tardigradepreservation. Many thanks to Lin Lee for ordering reagents, and supplies.

## Author Contributions

**SS, SR, AG-T, PDL** created crowdfunding project, raised funds, experimental design and execution, **SS, SR, PDL** performed data analysis and wrote draft. **AG-T, GMM** edited draft.

