## Supplemental Data for "A low-cost and easy-to-use cell preservation reagent for 4°C or room temperature sample storage"

### Supplemental Materials and Methods

**Cell Culture** Low passage primary human diploid foreskin fibroblast cells were used for preservation experiments. The cells were cultured in DMEM (Gibco, ThermoFisher Scientific) supplemented with 10% fetal bovine serum and 1% PenStrep (100 U/mL Penicillin /10 μg/mL Streptomycin). The cells were maintained under standard cultural conditions at 37°C in an atmosphere of 5% $CO_2$ and 5% $O_2$ in a Heracell CO2 incubator (ThermoFisher Scientific, Waltham, MA). Gas phase O2 tensions were controlled by continuous injection of appropriate amount of medical grade N2 to reach the target levels of oxygen . Cells were seeded at equal cell densities in 12-well, 6-well, or 60 mm dishes for the isolations of protein, DNA, or RNA .

**Nucleic Acids** DNA was extracted using in-house reagents based on the Gentra-Puregene methods (Qiagen, Germantown, MD). RNA was extracted using Trizol, as per the manufacture’s protocol. RNA and DNA concentrations were determined using a Nanodrop Lite Nano Spectrophotometer (ThermoFisher Scientific, Waltham, MA) and visualized on 1% agarose e-gel Power Snap (ThermoFisher Scientific, Waltham, MA).

**Preservation** Reagent #1 is PBS, pH 7.4, the base of all conditions. Reagent #2 is PBS plus 10 mM EDTA. Reagent #3 contains PBS, plus 10 mM EDTA and 50 mM Trehalose (ThermoFisher Scientific, Waltham, MA). Reagent #4A is PBS, plus 10 mM EDTA and 5% fetal bovine serum. Reagent #4B contains PBS, pH 7.4 plus 10 mM EDTA and 5% molecular grade BSA (ThermoFisher Scientific, Waltham, MA).

The volume of preservation reagent was different for each vessel size. 75 μl, 100 μl, and 150 μl of reagents 1-4, were used for 12-well, 6-well, and 60 mm dishes respectively. A 9-inch oscillating fan (SMC, China) was used to rapidly dry each dish of reagent and cells. The dry dishes of cells were immediately placed in a glass desiccator in a dark cabinet. Less than 10% humidity was achieved using drierite anhydrous calcium sulfate (W.A. Hammond Drierite Co Ltd, Xenia, OH) and food grade silica packs (Dry & Dry, Silicagel Factory, Brea, CA).

For each preservation trial, untreated samples of equal cell densities were frozen at

-80C for the same duration as the dry preservation samples. Samples for all preservation reagent conditions were harvested at 1 week, 1, 2, and 3 months. Fresh samples of equal cell densities were harvested on day zero to represent the starting levels of protein, DNA, and RNA .

**Supplemental Figures**

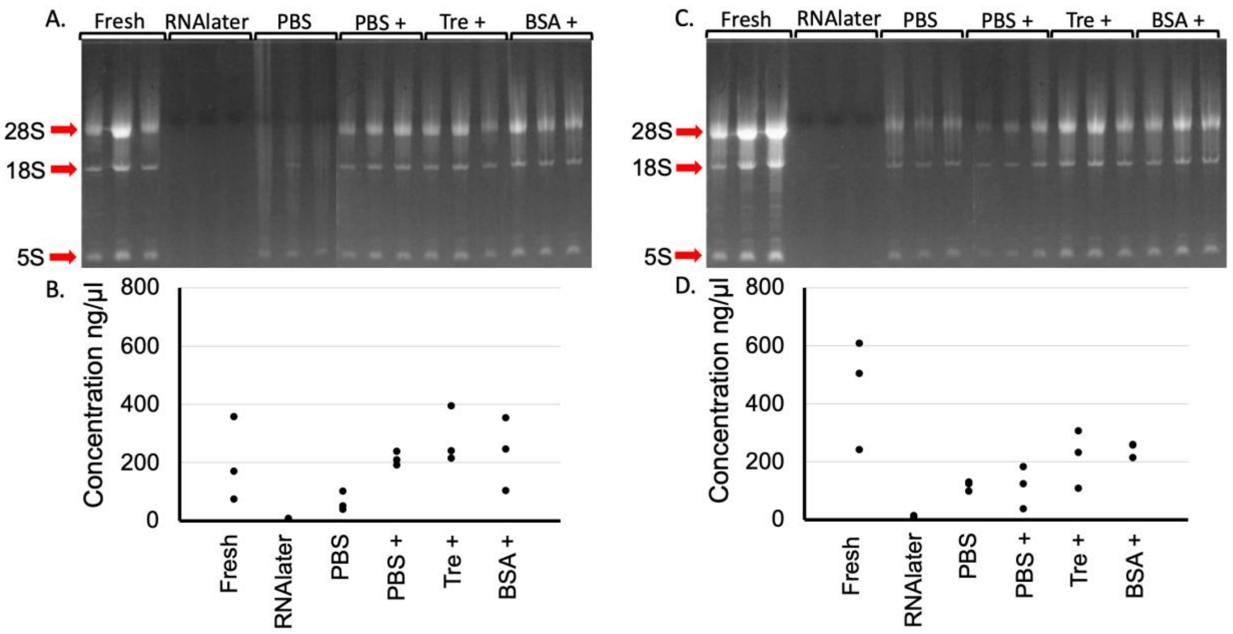


Supplemental Figure 1. Replicates of RNA extracted from cells stored in solution at 4℃ for one week. A) and C) Agarose gel with RNA from freshly harvested cells and compared with RNA from cells stored in RNAlater, PBS, PBS +, Tre +, and BSA +. Red arrows denote 28S, 18S, and 5S. Samples are shown as a composite of two gels resolved consecutively, reflecting all of the samples per cell density plating. B) and D) Graph depicting RNA concentrations in ng/µl. All samples in paired gel and graph reflect three replicate plates per preservation condition. N = 6

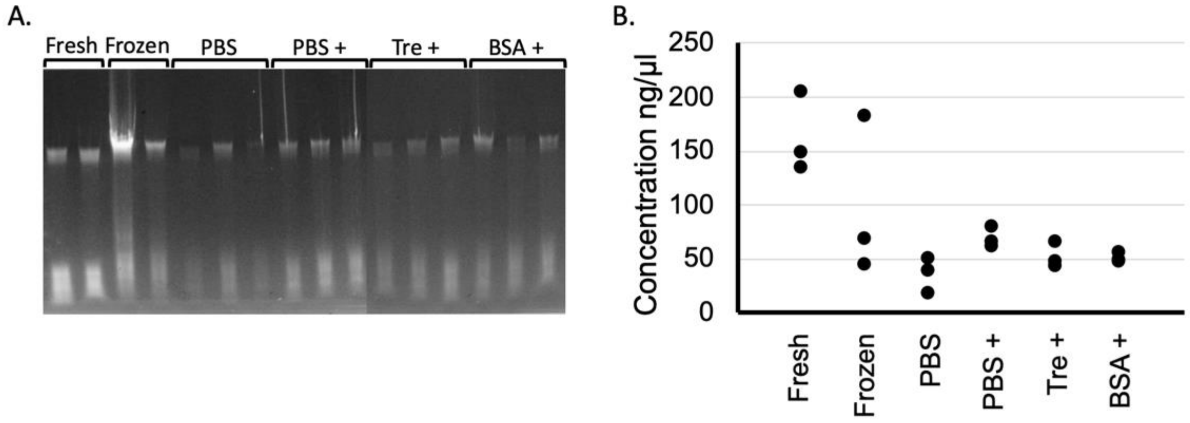


Supplemental Figure 2. DNA can be purified from A) Representative 1% agarose gel with DNA from freshly harvested cells, frozen, or stored in PBS, PBS +, Tre +, and BSA +. DNA gels are stained with SYBR safe. B) Scatter plot of DNA concentration for freshly harvested samples and samples stored for one week frozen at -80℃, or at 4℃ in PBS, PBS +, Tre +, or BSA +. Concentration unit is ng/µl.

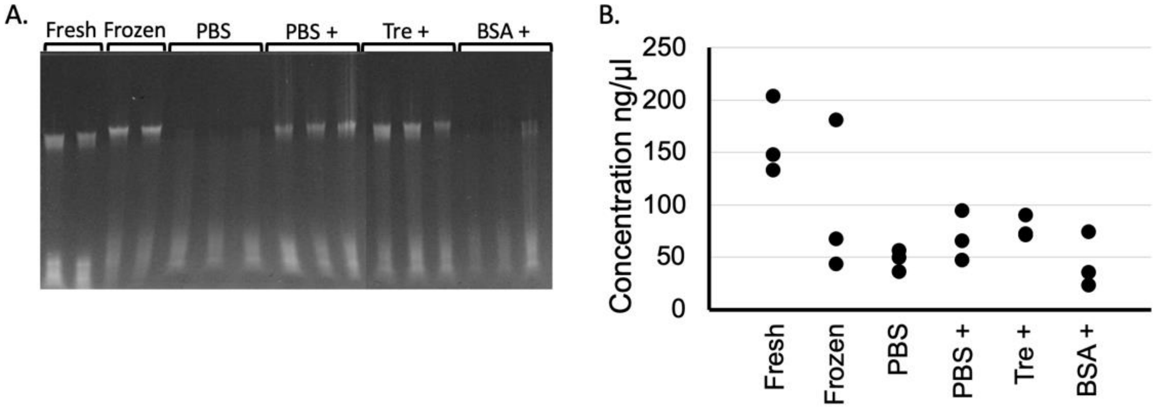


Supplemental Figure 3. DNA can be purified from samples that were stored dry in a desiccator for one week. A) Representative 1% agarose gel with DNA from freshly harvested cells, frozen, or stored in PBS, PBS +, Tre +, and BSA +. Samples are shown as a composite of two gels resolved consecutively, reflecting all of the samples per cell density plating. DNA gels are stained with SYBR safe. B) Scatter plot of DNA concentration for freshly harvested samples and samples stored for one week frozen at -80℃, or at 4℃ in PBS, PBS +, Tre +, or BSA +. Concentration unit is ng/µl
